# Double-digest RAD-sequencing: do wet and dry protocol parameters impact biological results?

**DOI:** 10.1101/445122

**Authors:** Tristan Cumer, Charles Pouchon, Frédéric Boyer, Glenn Yannic, Delphine Rioux, Aurélie Bonin, Thibaut Capblancq

## Abstract

1. Next-generation sequencing technologies have opened a new era of research in genomics. Among these, restriction enzyme-based techniques such as restriction-site associated DNA sequencing (RADseq) or double-digest RAD-sequencing (ddRADseq) are now widely used in many population genomics fields. From DNA sampling to SNP calling, both wet and dry protocols have been discussed in the literature to identify key parameters for an optimal loci reconstruction.
2. The impact of these parameters on downstream analyses and biological results drawn from RADseq or ddRADseq data has however not been fully explored yet. In this study, we tackled this issue by investigating the effects of ddRADseq laboratory (*i.e.* wet protocol) and bioinformatics (*i.e.* dry protocol) settings on loci reconstruction and inferred biological signal at two evolutionary scale using two systems: a complex of butterfly species (*Coenonympha sp.*) and populations of Common beech (*Fagus sylvatica*).
3. Results suggest an impact of wet protocol parameters (DNA quantity, number of PCR cycles during library preparation) on the number of recovered reads and SNPs, the number of unique alleles and individual heterozygosity. We also found that bioinformatic settings (*i.e.* clustering and minimum coverage thresholds) impact loci reconstruction (*e.g.* number of loci, mean coverage) and SNP calling (*e.g.* number of SNPs, heterozygosity). We however do not detect an impact of parameter settings on three types of analysis performed with ddRADseq data: measure of genetic differentiation, estimation of individual admixture, and demographic inferences. In addition, our work demonstrates the high reproducibility and low rate of genotyping inconsistencies of the ddRADseq protocol.
4. Thus, our study highlights the impact of wet parameters on ddRADseq protocol with strong consequences on experimental success and biological conclusions. Dry parameters affects loci reconstruction and descriptive statistics but not biological conclusion for the two studied systems. Overall, this study illustrates, with others, the relevance of ddRADseq for population and evolutionary genomics at the inter- or intraspecific scales.

## INTRODUCTION

For a decade, next-generation sequencing (NGS) technologies have opened a new era in the large field of molecular ecology In particular, the advances in sequencing capabilities have deeply changed the field of population genetics, by providing tremendous amount of sequence data/information (10 to 100 thousand markers) at a relatively low cost. (Andrews et al., 2014; da Fonseca et al., 2016). Whole genome re-sequencing (WGR) methods, providing the highest marker density among the current genomic methods, notably appear very useful to investigate many questions in evolutionary biology and ecology (Fuentes-Pardo & Ruzzante, 2017). WGR has however a limited relevance for non-model species, because a reference genome is not always available and because it requires considerable sequencing and computing efforts (Fuentes-Pardo & Ruzzante, 2017). Stemming from these limitations, reduced-representation sequencing methods have been developed. These approaches include restriction-site associated DNA sequencing (RAD-sequencing), sequencing of transcribed DNA from mRNA (RNA-sequencing), and whole-exome sequencing (WES). Overall, reduced-representation sequencing methods allow accessing numerous homologous loci with a great taxa coverage at a relatively low cost (Fuentes-Pardo & Ruzzante, 2017). Among these methods, RAD-sequencing, or RADseq (Miller, Dunham, Amores, Cresko, & Johnson, 2007; Baird et al., 2008), is certainly the most popular method to obtain thousands of single nucleotide polymorphisms (SNPs) for non-model species (K. R. Andrews, Good, Miller, Luikart, & Hohenlohe, 2016). The principle of RADseq is to use restriction enzymes to subsample the genome of multiple individuals at homologous genomic locations (Miller et al., 2007; Baird et al., 2008). The resulting DNA fragments are then sequenced and compared among individuals to detect SNPs. Since its origin, this technique has been transformed into a variety of related approaches (Peterson, Weber, Kay, Fisher, & Hoekstra, 2012; S. Wang, Meyer, Mckay, & Matz, 2012; Toonen et al., 2013; Campbell, Brunet, Dupuis, & Sperling, 2018). Among these, double-digest RADseq, or ddRADseq (Peterson et al., 2012), is highly customizable as regards the final number of loci, depending on the choice of enzymes and range of fragment size selected. The ddRADseq approach has been applied with success to many purposes including population genetic studies (Kjeldsen et al., 2016; Black, Seears, Hollenbeck, & Samollow, 2017; Sherpa, Rioux, Goindin, et al., 2018), phylogenetic reconstructions (DaCosta & Sorenson, 2016; Vargas, Ortiz, & Simpson, 2017; Boubli et al., 2018; Lee et al., 2018; Sherpa, Rioux, Pougnet-Lagarde, & Després, 2018), demographic inferences (Capblancq, Després, Rioux, & Mavárez, 2015; Nunziata, Lance, Scott, Lemmon, & Weisrock, 2017; Settepani et al., 2017; Elleouet & Aitken, 2018) and landscape genetic analyses (Saenz-Agudelo et al., 2015; Johnson, Gaddis, Cairns, Konganti, & Krutovsky, 2017). Despite the recognized advantages of the ddRADseq technique, several limitations and weaknesses arose in the literature (Davey et al., 2013; K. R. Andrews et al., 2016; Lowry et al., 2017). The main concerns are related to both the wet laboratory and bioinformatics procedures associated with the method (Puritz et al., 2014; Mastretta-Yanes et al., 2015; Shafer et al., 2017).

The frequency and distribution of restriction sites in the genome vary considerably depending on the species and the pair of enzymes considered (Herrera, Reyes-Herrera, & Shank, 2015). Among wet lab specific aspects, the choice of enzymes is therefore critical for appropriate genome subsampling through a ddRADseq procedure. For example, this choice will influence the number of digested fragments, their location in the genome, and their size distribution (Burns et al., 2017; Y. Wang et al., 2017). DNA quality also influences SNPs recovery, because degraded (*i.e.* fragmented) DNA can greatly lower the efficiency of restriction enzyme-based techniques, by inducing a loss of recovered fragments (Graham et al., 2015). Amplification of ddRADseq or RADseq fragments during library preparation has also been pointed out as a potential critical step (Davey et al., 2013; Mastretta-Yanes et al., 2015). Indeed, non-homogeneous amplification of RAD fragments can lead to a substantial loss of alleles due to unbalanced RAD fragments coverage (Andrews & Luikart, 2014; Andrews et al., 2014; Puritz et al., 2014). Furthermore, the number of PCR cycles during library preparation is generally low to minimize PCR artifacts (such as PCR errors) in the RAD tags sequences (Hohenlohe, Catchen, & Cresko, 2012; Peterson et al., 2012).

Another important concern about RAD-based methods is the bioinformatic treatment of sequences and the reconstruction of RADseq loci (*e.g.* Shafer et al., 2017). The principle of the RADseq technique relies on the identification of homologous loci among individuals. This task is implemented by clustering single-copy loci according to a similarity threshold, which is determined using either distance-based (*e.g. STACKS*; Catchen, Hohenlohe, Bassham, Amores, & Cresko, 2013), or global alignment (*e.g. pyRAD*; Eaton, 2014) methods. In both cases, stringent parameter settings will avoid the clustering of paralogs but can also split highly divergent single-copy loci in different clusters (Catchen et al., 2013; Eaton, 2014). Coverage is another important parameter for loci reconstruction. A minimum number of reads is generally set in order to take into account an allele or not (Catchen et al., 2013). Defining a high threshold value can induce a loss of alleles via insufficient coverage, while a too low value will not discard rare sequences originating from PCR or sequencing errors (Paris, Stevens, & Catchen, 2017). The influence of parameter settings on quality and quantity of recovered fragments and SNPs has therefore been widely tested (Eaton, 2014; Mastretta-Yanes et al., 2015; Paris et al., 2017; Rochette & Catchen, 2017; Shafer et al., 2017) and it sometimes impacts downstream population genomics analyses (Ilut, Nydam, & Hare, 2014; Harvey et al., 2015; Willis, Hollenbeck, Puritz, Gold, & Portnoy, 2017).

Finally, the impact of missing data, which are inherent to any genotyping technique, has also been evaluated over the years (Arnold, Corbett-Detig, Hartl, & Bomblies, 2013; Gautier et al., 2013; Malinsky, Trucchi, Lawson, & Falush, 2018). Missing data can be due to some extent to an experimental lack of reproducibility, but more frequently to polymorphism in restriction sites. This polymorphism leads to allele drop-out (ADO) for the individuals lacking the restriction site in one or two of the homologous chromosomes. ADO directly influences the estimation of genetic variation and diversity (Davey et al., 2013; Gautier et al., 2013; Cariou, Duret, & Charlat, 2016). It has been particularly investigated in phylogenetic studies (Cariou, Duret, & Charlat, 2013; Eaton, 2014; DaCosta & Sorenson, 2016) because of the direct correlation between ADO and the divergence time among lineages and species (Cariou et al., 2013).

The scientific community has accumulated expertise about RAD-based methods over the last decade to make better use of such techniques while some critical issues deserve further investigations. Indeed, if the proximal consequences have been investigated (*e.g.* number of SNPs or deficit in heterozygosity), the distal consequences (effect on genetic structure estimation, demographic inferences, etc.) remain rarely explored in most cases. The impact of the minimum coverage or the similarity threshold for sequences clustering has for example already been largely discussed in the literature. However, if the community accepts that these parameters directly influence the number and coverage of loci or the percentage of missing data in the final genetic matrix (Catchen et al., 2013), their potential effects on the biological signal unraveled by downstream analyses are not systematically tested and this issue lacks empirical investigation (but see Mastretta-Yanes et al., 2015; Rodríguez-Ezpeleta et al., 2016; Shafer et al., 2017). In addition, some of the wet laboratory procedures are thought to be critical for the success of the experiment (Peterson et al., 2012; Mastretta-Yanes et al., 2015), like the initial DNA quantity and number of PCR cycles, but they have never been experimentally evaluated.

This study aims to examine these steps of the ddRADseq procedure, providing a novel contribution to the literature, in an animal system at an interspecific level (in the butterfly species complex of *Coenonympha*) and in plants at an intraspecific scale (in tree populations of European/common beech, *Fagus sylvatica*). We first evaluate the impact of both initial DNA quantity and number of PCR cycles on the experiment results, and the reproducibility of our wet protocol by evaluating the percentage of genotyping inconsistencies and missing fragments between replicates. Concerning the bioinformatic treatment, we investigate the influence of the minimum coverage and the similarity threshold during the loci reconstruction on three types of analyses based on ddRADseq data: genetic differentiation (evaluated using *F*_ST_ estimation and Principal Component Analysis), genetic structure (genetic clustering) and demographic inferences (Approximate Bayesian Computation method).

## MATERIALS AND METHODS

### Sampling

This study is based on a total of 108 samples, including 58 individuals of *Fagus sylvatica* (deciduous tree; genome size around 600 Mbp) and 50 individuals of *Coenonympha* sp. (three butterfly species: *C. arcania*, *C. gardetta* and *C. macromma*; genome size around 300 Mbp). Depending on the conditions and settings tested (Fig. 1), different numbers of individuals and populations were used (see Table S1).

**Figure 1:**
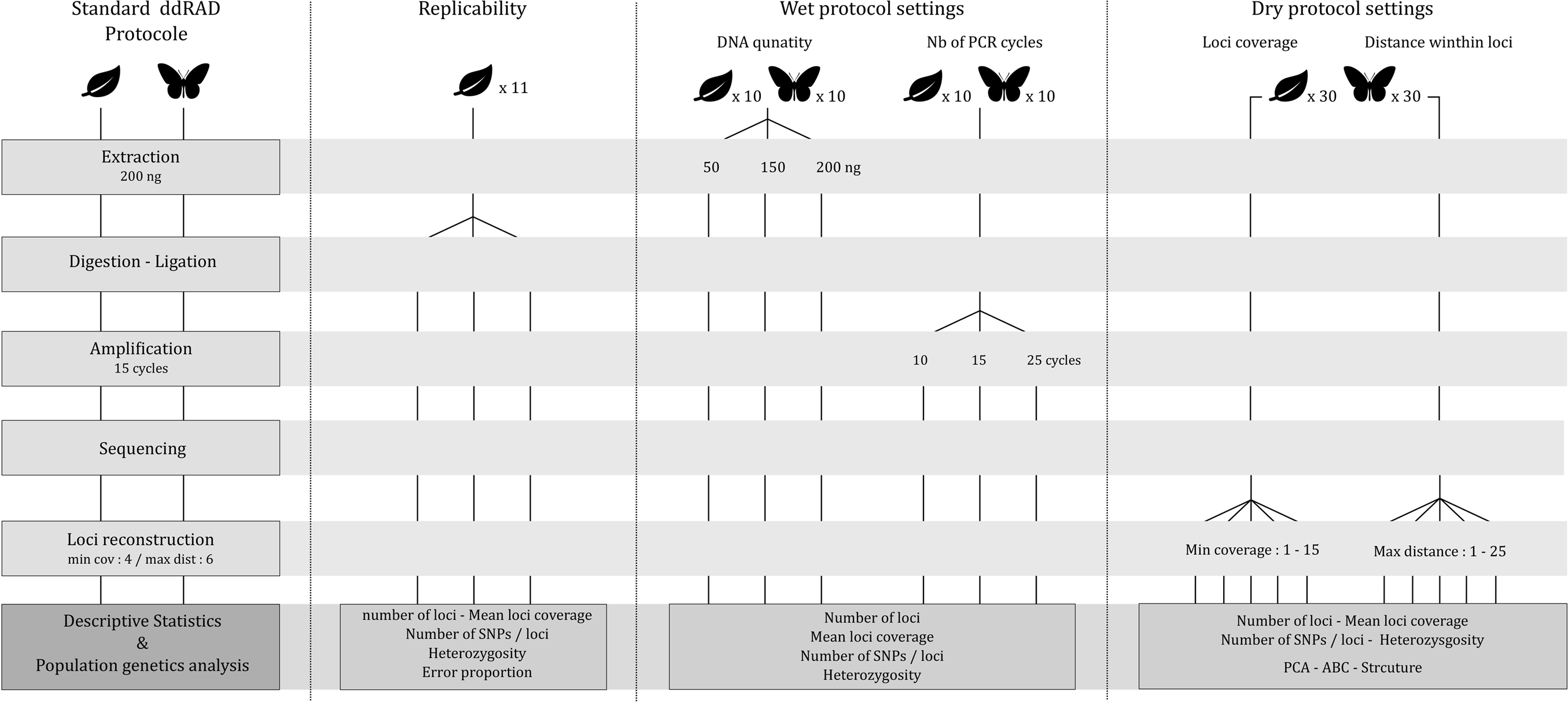
Overview of the experimental design used in this study.

### Standard ddRADseq protocol

A double-digested RAD experiment was conducted on individuals using a common protocol for both the wet and dry parts of the procedure. The protocol was the same for all samples, except for some parameter settings as described in the following sections: “*Setting tests for the wet laboratory protocol*” and “*Tests on bioinformatics parameters*”.

#### Library preparation

DNA was extracted from one leaf for *Fagus sylvatica* samples and from the complete thorax of individual *Coenonympha* butterflies using a DNeasy Blood & Tissue Kit (QIAgen) following manufacturer’s instructions. For each individual, 200 ng of genomic DNA were double-digested with 10 units of each *Pst*I and *Msp*I (New England Biolabs Inc.) at 37°C for two hours in a final volume of 34μL, using the CutSmart buffer provided with the enzymes. Digestion was further continued with ligation of the P1 (with individual tags) and P2 adapters (see Peterson et al., 2012) by adding to each sample 10 units of T4 DNA ligase (New England Biolabs Inc.), adapters P1 and P2 in 10-fold excess (compared to the estimated number of restriction fragments), 1μL of 10mM ribo-ATP (New England Biolabs Inc.) and once again two units of *PstI* and *MspI* enzymes. This simultaneous digestion-ligation reaction was performed on a thermocycler using 60 cycles of a succession of 2 min at 37°C for digestion and 4 min at 16°C for ligation. An equal volume of all the digested-ligated fragment mixtures were pooled and purified using magnetic beads (Agencourt AMPure XP of Beckman Coulter, or NucleoMag of Macherey Nagel) with a DNA/beads ratio equal to 1/1.5. Fragments were size-selected in a range between 250 and 500 bp on agarose gel (1.6%) and excised bands purified with the QIAquick Gel Extraction Kit (Qiagen). The ddRADseq library obtained was amplified independently eight times by PCR, and the obtained PCR products were then pooled, in order to minimize the impact of potential PCR errors. We used the following PCR mix: a final volume of 20 μL containing 1 μL of DNA template, 10 mM of dNTPs, 10 μM of each PCR primer (Peterson et al., 2012) and 2U/μL of *Taq* Phusion-HF (New England Biolabs Inc.); and the following PCR program: an initial denaturation at 98°C for 30 seconds; 15 cycles of 98°C for 10 sec, 66°C for 30 sec and 72°C for 1 min; followed by a final extension at 72°C for 10 min. The amplified ddRADseq library was purified with magnetic beads and sequenced on half a lane of an Illumina Hi-Seq 2500 sequencer (paired-end 2 × 125 bp, Fasteris SA).

#### Bioinformatic treatment

DNA sequences of *Fagus sylvatica* and *Coenonympha* sp. libraries were used to call SNP genotypes (total: ~100 million reads). We developed a homemade pipeline called ProcessMyRAD (fully available at https://github.com/cumtr/pmr) to automatically perform the different steps leading from the raw reads to genotype data. To call the genotypes, ProcessMyRAD relies on the *STACKS* pipeline (Catchen et al., 2013). To reconstruct loci, the *STACKS* procedure needs to set three thresholds: the minimum number of reads to consider an allele (**m**), the maximum number of mismatches allowed between two alleles to reconstruct a locus (**M**), and the maximum number of mismatches allowed between two individual loci to consider them as homologous (**n**).

### Setting tests for the wet laboratory protocol

#### Impact of initial DNA amount and number of PCR cycles

We evaluated the impact of two parameters on the experiment results: 1) the DNA quantity used for the initial digestion/ligation step; and 2) the number of PCR cycles used to produce the final library.

For 10 samples of *Coenonympha* and 10 samples of *Fagus sylvatica* (Table S1), we repeated the ddRADseq lab experiment three times for each sample with the standard protocol described above but with different quantities of DNA during the first step: 50, 150 or 250 ng of initial genomic DNA. Similarly, we used 10 samples of *Coenonympha* sp. and 10 samples of *Fagus sylvatica* to repeat three times the ddRADseq lab experiment, with different numbers of PCR cycles in the final step of the protocol: 10, 15 or 25 cycles. We then sequenced the resulting libraries all together.

The sequences resulting from these tests were treated with the *STACKS* program and the following clustering parameters: **m**=4, **M**=6, **n**=8 (based on the results of the section “Bioinformatics tests” for **m** and **M**, and with **n**=**M**+2 to increase the number if inter-individual matches), keeping only one SNP by ddRADseq fragment. To estimate the impact of the wet lab treatment on alleles frequencies, we did not filter out alleles based on their frequency. On the resulting genetic datasets, we determined the number of polymorphic fragments, the mean fragment coverage, the number of SNPs in the fragments, the individual heterozygosity, and the proportion of private alleles in individuals.

#### Experimental reproducibility

We assessed the reproducibility of our laboratory protocol by repeating the experiment, since the digestion/ligation step, three times for 11 *Fagus sylvatica* individuals. Each replicate of these triplets was processed with the same protocol and was sequenced within the same Illumina sequencing run. All sequences obtained were treated together with the ProcessMyRAD pipeline with **m**=4, **M**=6, **n**=8 and a minor allele frequency of 0.1 (corresponding to at least three individuals to keep the allele). A locus was kept only if sequenced in at least 50% of the 33 replicates.

For each replicate, the number of ddRADseq fragments, the mean fragment coverage, the proportion of polymorphic fragments and the individual heterozygosity were estimated. These different parameters were then compared among replicates to assess the intra and inter-replicate variability. We also evaluated reproducibility by performing a Principal Component Analysis on genetic data with the R-package *adegenet* (Jombart, 2008), and looking at the distances between replicates in the PCA projection space. In addition, the replicates were used to estimate the proportion of inconsistencies in our final genetic dataset. These inconsistencies can take two different forms: errors of genotypes (due to PCR errors or ADO) or fragment absence (due to a lack of reproducibility of the experiment). We measured the genotyping inconsistency rate by identifying the proportion of loci with inconsistencies among the three replicates. Then, by looking at the fragments in the three replicates, we could estimate the proportion of “true” fragment absences, when the fragment was missing in all three replicates, and the proportion of “false” fragment absences, when the fragment was missing in just one or two of the replicates.

### Test on bioinformatic parameters

#### Impact of bioinformatic treatment

We estimated the influence of the **m** (*ustacks*) and **M** (*cstacks*) values on ddRADseq fragment reconstruction and downstream analyses.

For this purpose, we used the standard ddRADseq wet protocol described above on 30 individuals of *Fagus sylvatica* coming from three different populations (Sainte Beaume, Digne, and Bauges, France; see Table S1) and 30 individuals of *Coenonympha* butterflies from three different species (*C. arcania, C. gardetta* and *C. macromma*; see Table S1). We then repeated the bioinformatic pipeline with different **m** values ranging from 1 to 15 and different values of **M** ranging from 1 to 25. When the **m** value varied, **M** and **n** were fixed to 6. When the **M** value varied, **m** was fixed to 4 and **n** was equal to **M**. For all these tests, the remaining steps of the procedure were exactly the same and the last step of genetic dataset export was performed by keeping only one SNP by RAD tag fragment and without any filtering on allelic frequency.

To estimate the influence of the **m** and **M** values on RAD tag fragment recovery, we determined the number of reconstructed fragments, their mean coverage and the proportion of polymorphic fragments for each value of **M** and **m** tested. We also evaluated the impact of these parameters on population genetics results by performing, for all **m** and **M** values, some of the most commonly used analyses using ddRADseq data (Capblancq et al., 2015; Kjeldsen et al., 2016; Black et al., 2017; Nunziata et al., 2017; Settepani et al., 2017; Elleouet & Aitken, 2018; Sherpa, Rioux, Pougnet-Lagarde, et al., 2018), i.e. mean individual heterozygosity, *F*_ST_ among populations (estimated with the *adegenet* R package (Jombart, 2008)), Principal Component Analysis (PCA, using the *adegenet* R package (Jombart, 2008)), genetic structure with sNMF (using the *LEA* R package (Frichot & François, 2015)) and evolutionary history reconstruction using Approximate Bayesian Computation (performed with the diyABC program (Cornuet et al., 2014)). The results of these analyses were then compared across the **m** and **M** ranges and with results from other population genetic studies on the same species *Coenonympha* sp. In Capblancq et al., 2015, and *Fagus sylvatica* in Capblancq *et al*. (in review).

## RESULTS

### Influence of DNA quantity and number of PCR cycles

#### DNA quantity

For all initial DNA quantity conditions, the library produced a mean of 3,172 fragments for *Coenonympha,* with a mean coverage of 24.6 reads per fragment, and 450 fragments for *Fagus sylvatica,* with a mean coverage of 19.5 reads per fragment. With 50 ng or 150 ng of genomic DNA as template, similar numbers of fragments and SNPs were recovered (around 3,500 fragments for *Coenonympha* and around 500 for *Fagus sylvatica*). If fragment coverage varies (from 15 to 30 for *Coenonympha* and from 15 to 23 for *Fagus sylvatica*), this does not have much impact on individual heterozygosity (He~ 0.125 for *Coenonympha* and He~0.25 for *Fagus sylvatica*). Conversely, we noticed that using 250 ng of DNA during the initial step of digestion/ligation could dramatically decrease fragment recovery (divided by 1.5) and SNPs identification for some individuals (Fig. 2). Using 250 ng of DNA also induced a greater variability among tested individuals (Fig. 2).

**Figure 2:**
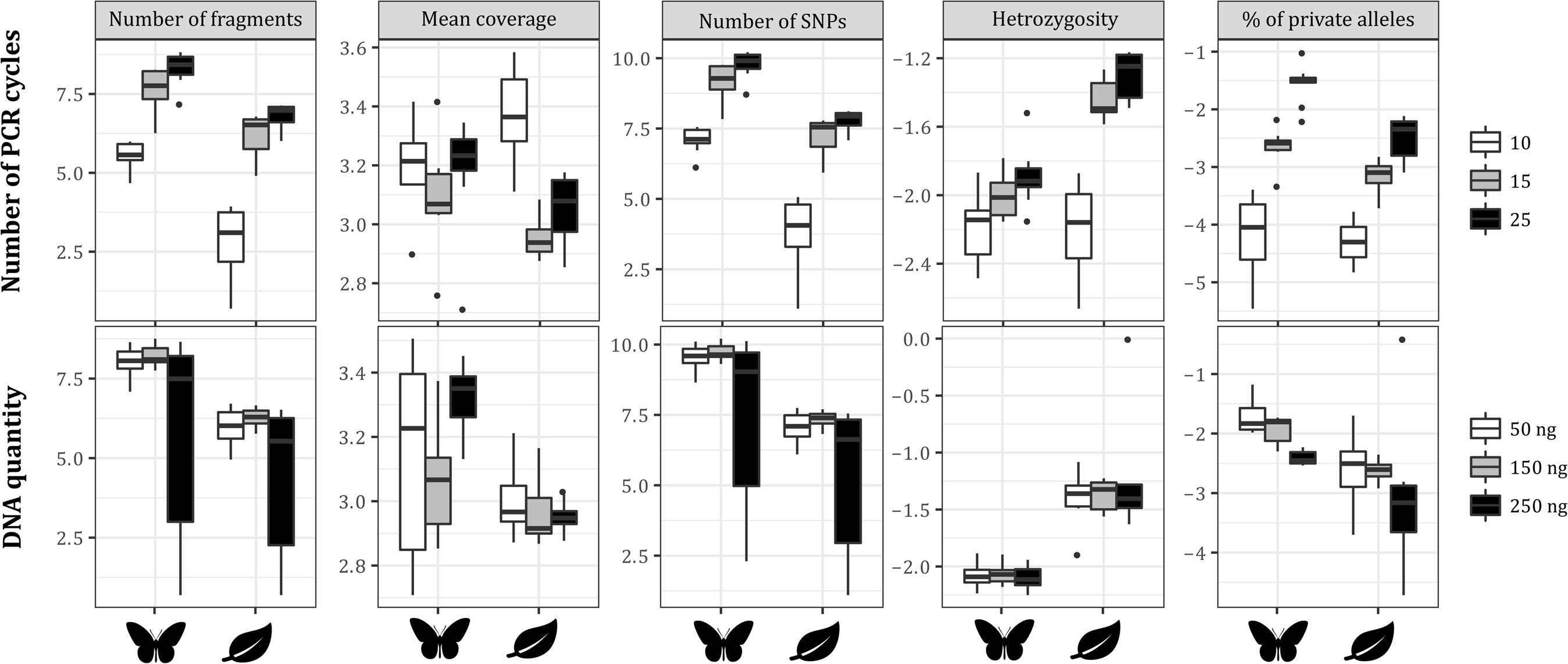
Impacts of initial DNA quantity (bottom) and number of PCR cycles (top) on ddRADseq results. For each condition and each biological model (*Coenonympha* sp. or *Fagus sylvatica*), boxplots present the number of recovered fragments, the mean coverage of these fragments, the number of identified SNPs, the individual heterozygosity and the number of private alleles in individuals. A log-transformation was performed on the results in order to simplify the comparison of the two models.

#### Number of PCR cycles

For all PCR conditions, the library produced a mean of 2,480 fragments for *Coenonympha* with a mean coverage of 23.9 reads per fragment and 520 fragments for *Fagus sylvatica* with a mean coverage of 23.3 reads per fragment. Again, the results showed a great variability depending on the PCR settings. Increasing the number of PCR cycles in the final library preparation had a positive effect on the number of fragments and SNPs recovery. The mean number of fragments for *Fagus sylvatica* ranged from 41 for 10 cycles to 1,028 for 25 cycles. Similar results were obtained for *Coenonympha* sp. for which the number of fragments varied from 350 to 4,500. The number of PCR cycles was also directly correlated with individual heterozygosity and with the number of private alleles in the individuals. For example, the individual heterozygosity increased from 0.09 to 0.27 for *Fagus sylvatica* individuals when the number of cycles increased from 10 to 25. In the same way, the number of private alleles doubled when the number of PCR cycles increased from 10 to 25 for both *Coenonympha* sp. and *Fagus sylvatica* samples. Finally, 10 cycles of PCR lowered Substantially the number of fragments and SNPs as well as the number of private alleles in the final genetic dataset.

### Reproducibility of the experiment and estimation of inconsistencies

The library of the 11 *Fagus sylvatica* triplicates produced a mean of 7,547 fragments with a mean coverage of 20.3 reads per fragments. The PCA performed on the complete genetic dataset showed very consistent results across the 11 tested individuals (Fig. 3). Inter-individual genetic variability was higher than inter-replicate genetic variability. All triplicates clustered in the PCA plot and the different individuals could easily be differentiated. Moreover, considering the eigenvalues, the 10 first axes retained most of the genetic variance within the three replicates * 11 sample tests (92%). For example, PC1 strongly discriminates individual VTX_H_83 from the rest of the sampling and PC2 differentiates individuals SB_H_42 and BG_1_1. This suggests that each PC captured parts of inter-individual genetic variability differentiating a particular individual from the remaining samples. Replicates did not seem to add any substantial genetic variability that could have been caught by the PCA.

**Figure 3:**
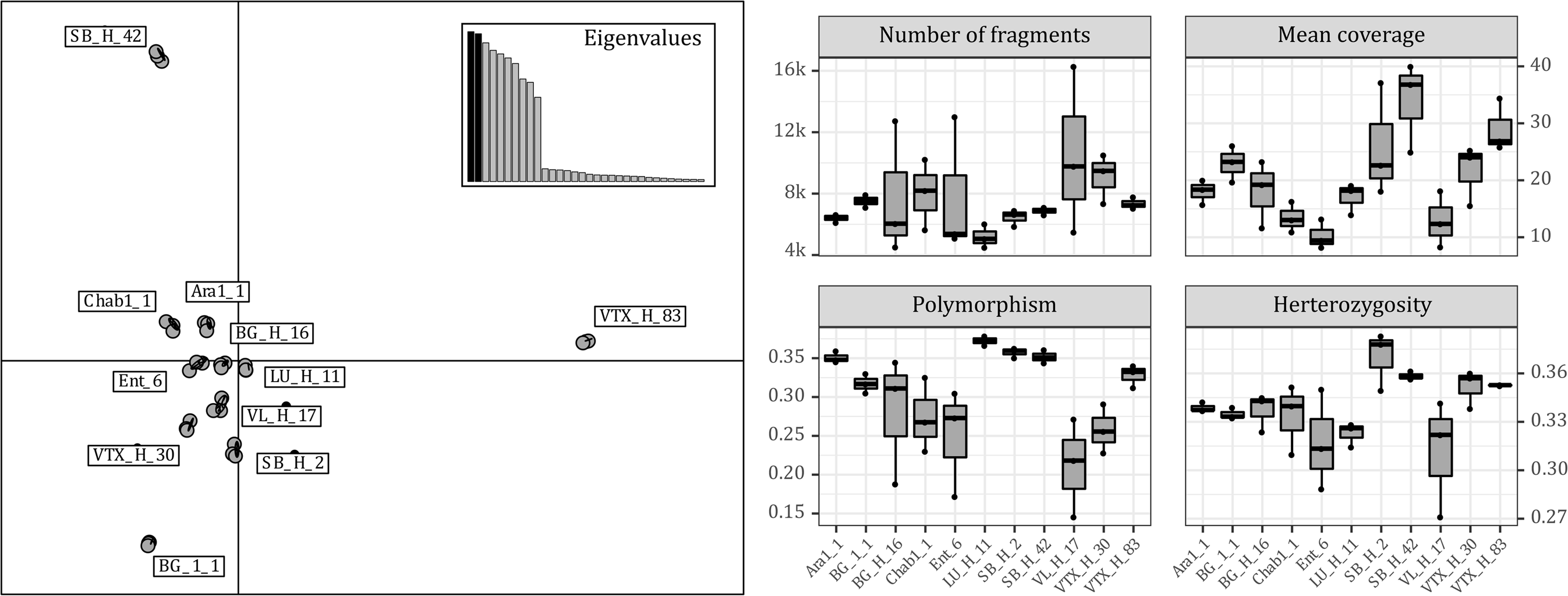
Reproducibility of the experimental wet protocol. The PCA on the left shows the inter-replicate genetic variability in comparison with inter-individual variability for *Fagus sylvatica* individuals. Each three-replicate group is circled by an ellipse. Boxplots on the right show the variation of the number of fragments, the mean coverage of the fragments, the proportion of polymorphic fragments and the individual heterozygosity within the three replicates of each individual.

Across all replicates and individuals, the number of recovered fragments varied from around 4,000 to 16,000; the mean coverage from 8 to 40; the proportion of polymorphic loci from 0.15 to 0.38, and the individual heterozygosity from 0.27 to 0.37 (Fig. 3). For seven individuals, the replicates returned almost exactly the same number of fragments, the same ratio of polymorphic loci and the same individual heterozygosity. Four individuals showed more contrasted results but with similar patterns across the different parameters we measured. No association between the initial DNA concentration after extraction and the consistency of ddRADseq results was observed (data not shown).

Regarding the estimation of genotype inconsistencies, the results were congruent across the 11 tested individuals (Fig. 4). The maximum inconsistency rate was just above 4% and the minimum is around 1.6%. Similarly, we obtained a good proportion of fragments recovery among replicates. Between 66% and 90% of the ddRADseq fragments were found in all three replicates (Fig. 4). The individuals with low fragment recovery rate were the exact same ones that showed a great variability in the reproducibility experiment (see above). Some fragments were missing for all three replicates, and the proportions of missing fragments were pretty homogeneous across individuals, varying from 5% to 13%. Finally, for all samples, a fair proportion of fragments (2 to 24%) was found in only one or two replicates, giving an estimation of fragment loss not due to restriction site polymorphism across individuals but to incomplete digestion, ligation, amplification or sequencing of these fragments.

**Figure 4:**
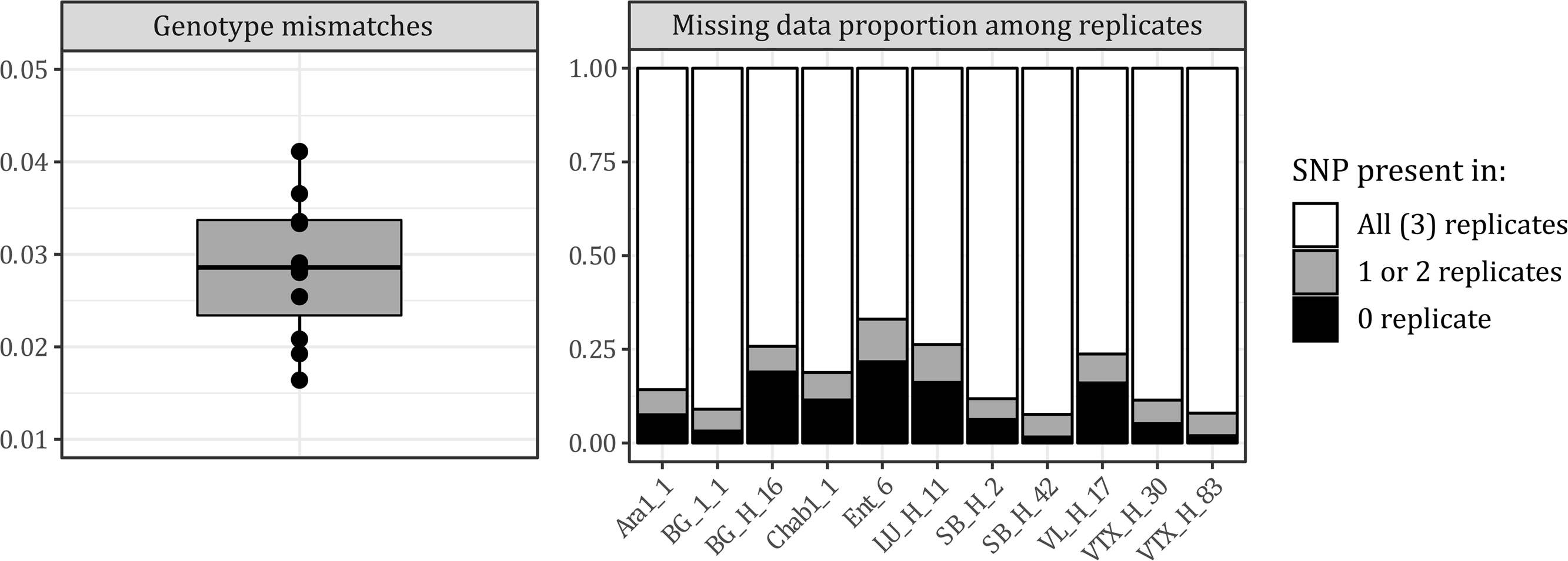
Left: boxplot of the genotyping inconsistency rate within *Fagus sylvatica* individuals. Right: proportion of loci found either in all the replicates (white), in only one or two replicates (grey) or in none of the three replicates (black).

### Influence of bioinformatic thresholds on the biological results

The ddRADseq libraries used for bioinformatic tests produced very variable numbers of fragments and coverage depending on the thresholds used along the analysis pipeline. With the parameter values **m** = 4 and **M** = 6, we obtained a mean of 3,246 fragments for *Coenonympha* individuals with a mean coverage of 40.16 reads per fragment and 11,018 fragments for *Fagus sylvatica* samples with a mean coverage of 25.18 reads per fragment.

The minimum coverage required to create a RAD tag locus during the first step of *STACKS* procedure (**m**) had a direct influence in the number of fragments, the mean coverage of the fragments and the number of SNPs identified in these fragments (Fig. 5). Furthermore, the pattern was very similar for the *Coenonympha* sp. and *Fagus sylvatica* models. An increase in **m** value was associated with a decrease in fragment recovery, the variation being particularly important between **m** = 1 and **m** = 2. Similarly, an increase of **m** value was associated with an increase in mean fragment coverage, ranging from 10x for a **m** value of 1 to 58x for a **m** value of 15 in *Coenonympha* sp. and from 5x for a **m** value of 1 to 30x for a **m** value of 15 in *Fagus sylvatica.* The number of SNPs identified in the fragments was also strongly affected by variation of the **m** threshold: the highest number of SNPs was reached for **m** = 2 and the lowest number for **m** = 15. Regarding individual heterozygosity, there was a slight increase when **m** increased. Heterozygosity ranged from 0.29 for **m** = 1 to 0.36 for **m** = 15 for *Fagus sylvatica* individuals, and from 0.13 to 0.15 for *Coenonympha* sp. samples.

**Figure 5:**
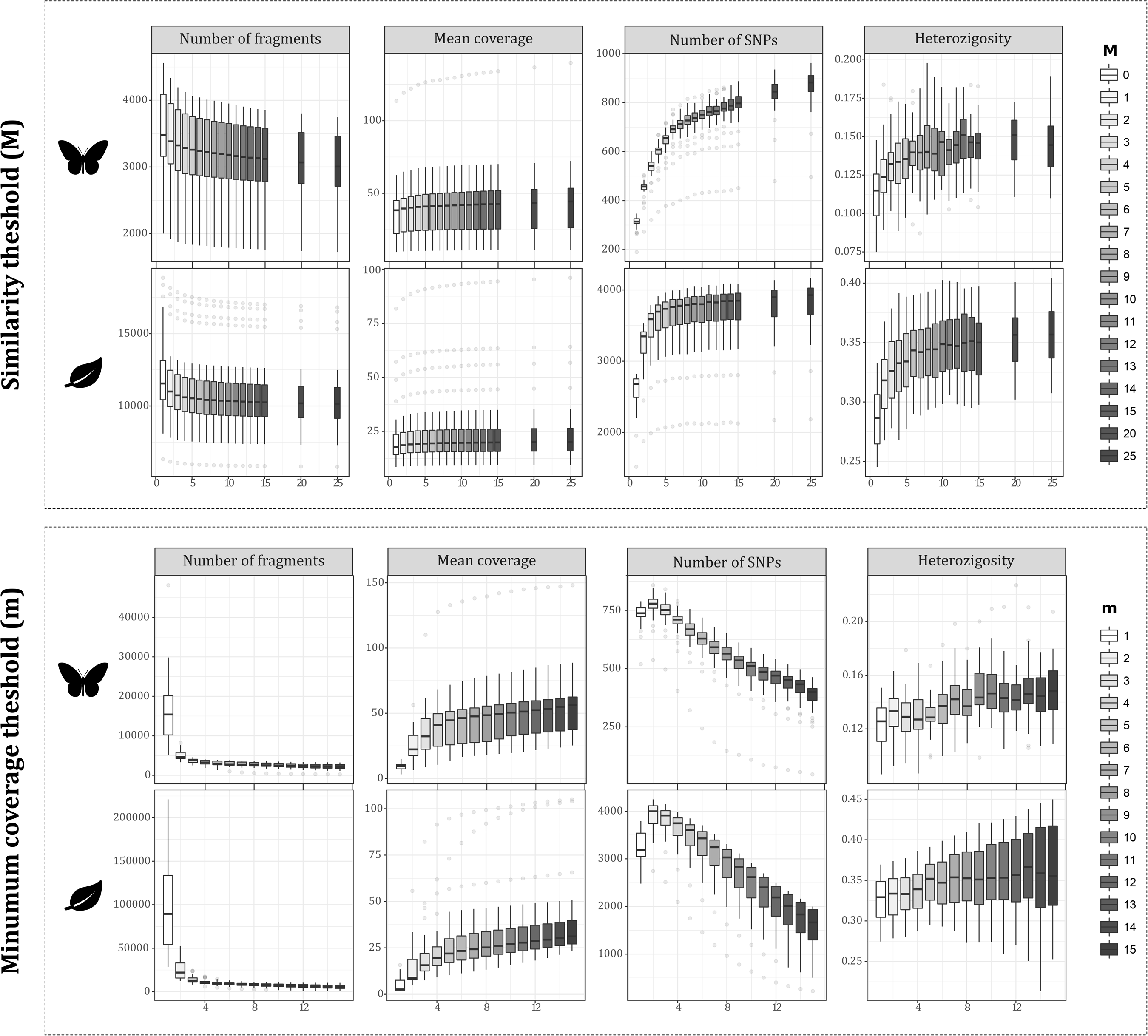
Impact of the *ustacks* thresholds **m** and **M** on the number of fragments, the mean coverage of the fragments, the number of SNPs, and the mean individual heterozygosity.

Nevertheless, the **m** parameter did not seem to influence any of the downstream population genetic analyses. No major difference among the **m** parameter settings was observed for *F_ST_* estimation among populations (Fig. 6), PCA and genetic clustering results (Fig. 6, Fig S1 and Fig. S2) or demographic inferences (Fig. 7). Here again, the results were similar for both the animal and plant models. While slight changes in *F_ST_* values or PCA scores were noticed when **m** varied, the populations remained differentiated in the same way and strength (Fig. 6). For example, the *F_ST_* values ranged from 0.28 to 0.33 between *Coenonympha arcania* and *C. gardetta* but the ranking of *F_ST_* values among the three pairs of species did not change depending on **m** (Fig. 6). An increasing **m** seemed to slightly influence the percentage of inertia retained by the first two PCs for both *Fagus sylvatica* and *Coenonympha* sp. (from 18 to 20% of the genetic variance, see Fig. 6) but population differentiation on the PCA was not affected. The Procrustes superimposition performed on the first two axes of the PCAs returns correlation coefficients superior to 0.96 between each pair of **m** values for *Coenonympha* and superior to 0.85 for *F. sylvatica* (Fig. S7). Regarding the genetic structure (sNMF analysis), the number of K selected with the cross-entropy criterion did not vary for *Coenonympha* samples and varied between 2, 3 and 4 for *Fagus sylvatica* individuals (Fig. 6). This variation was due to very close values of cross-entropy for K = 2, 3 and 4 (Fig. S2). Similarly, the differentiation of species groups in the sNMF analysis remained exactly the same across the range of **m** values (Fig. 6). The Procrustes superimposition performed on the percentage of assignation to the three main clusters obtained with sNMF returns correlation coefficients superior to 0.975 between each pair of **m** values for *Coenonympha* and superior to 0.99 for *F. sylvatica* (Fig. S8). Finally, we did not detect any influence of the **m** parameter on the estimations of population size, divergence time or hybridization rate through ABC procedure (Fig. 7). All model parameters showed approximately the same posterior distribution whatever the **m** value, with only a small variation between the maximum and the minimum of the estimates across the **m** range (Table S2).

**Figure 6:**
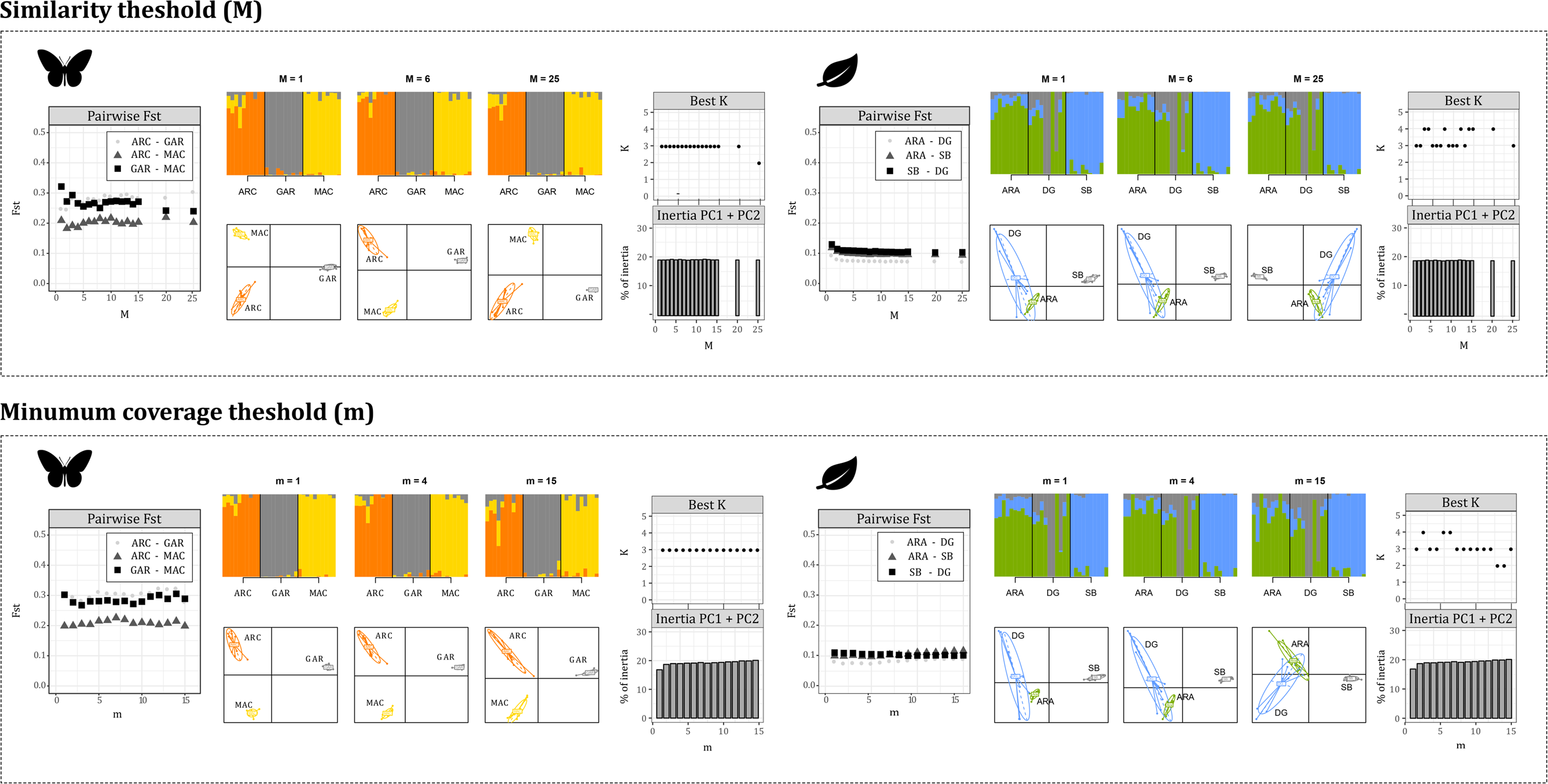
Impact of *ustacks* thresholds **m** and **M** in *F*_ST_ between pairs of populations, genetic differentiation and structure (PCA and sNMF results) for *Fagus sylvatica* and *Coenonympha sp.* individuals. Only the results for **m** =1, 4 and 15 and **M** = 1, 6 and 25 are shown (see Fig. S1 to S6 for the complete results). Ellipses in the PCA distinguish the different populations or species, sNMF results are shown for K = 3 which is the best number of clusters in almost all cases according to the cross-entropy criterion.

The maximum number of mismatches accepted between two stacks of sequences to merge two alleles in one locus (**M**) greatly influenced the number of recovered fragments, the number of identified SNPs and individual heterozygosity (Fig. 5). When **M** varied from 1 to 25 the number of recovered ddRADseq fragments decreased from 12,000 to 10,000 for *Fagus sylvatica* and from 3,500 to 3,000 for *Coenonympha* sp. On the opposite, the number of SNPs identified increased rapidly for the first **M** values of the range (1-6) until a plateau was reached around 3,800 fragments for *Fagus sylvatica* individuals. The influence of **M** on individual heterozygosity was clear for **M** values between 1 and 6, for which heterozygosity increased with **M**. For higher values of **M**, the relationship was less obvious and the variation of individual heterozygosity did not seem to follow the variation of **M**.

In agreement with the results obtained for the **m** parameter, population genetic analyses showed very consistent results across the range of tested **M** values. Again, no substantial effect was observed for *F_ST_* values among populations, PCA results, genetic clustering results or demographic inferences when **M** value varied (Fig. 6, Fig. 7, Fig. S4 and Fig. S5). **M** variation did not impact PCA, neither in terms of population differentiation, nor in terms of percentage of inertia of the two first axes. The Procrustes superimposition performed on the first two axes of the PCAs returns correlation coefficients superior to 0.95 between each pair of **m** values for *Coenonympha* and superior to 0.85 for *F. sylvatica* (Fig. S7). For genetic structure however, we observed a slight change in the number of K selected by the cross-entropy criterion. Here again, it is rather due to close cross-entropy values at K = 3 and K = 4 than to a real variation across the **M** range (Fig. S5). Even though, neither the genetic grouping of individuals nor the percentage of assignation varied across the range of **M** value (Fig. 6 and Fig. S8). Finally, all parameters inferred during the ABC analysis showed very consistent distributions depending on the **M** value used for the sequence clustering (Fig. 7), with only a small variation between the maximum and minimum of the estimates across the **M** range (Table S2).

**Figure 7:**
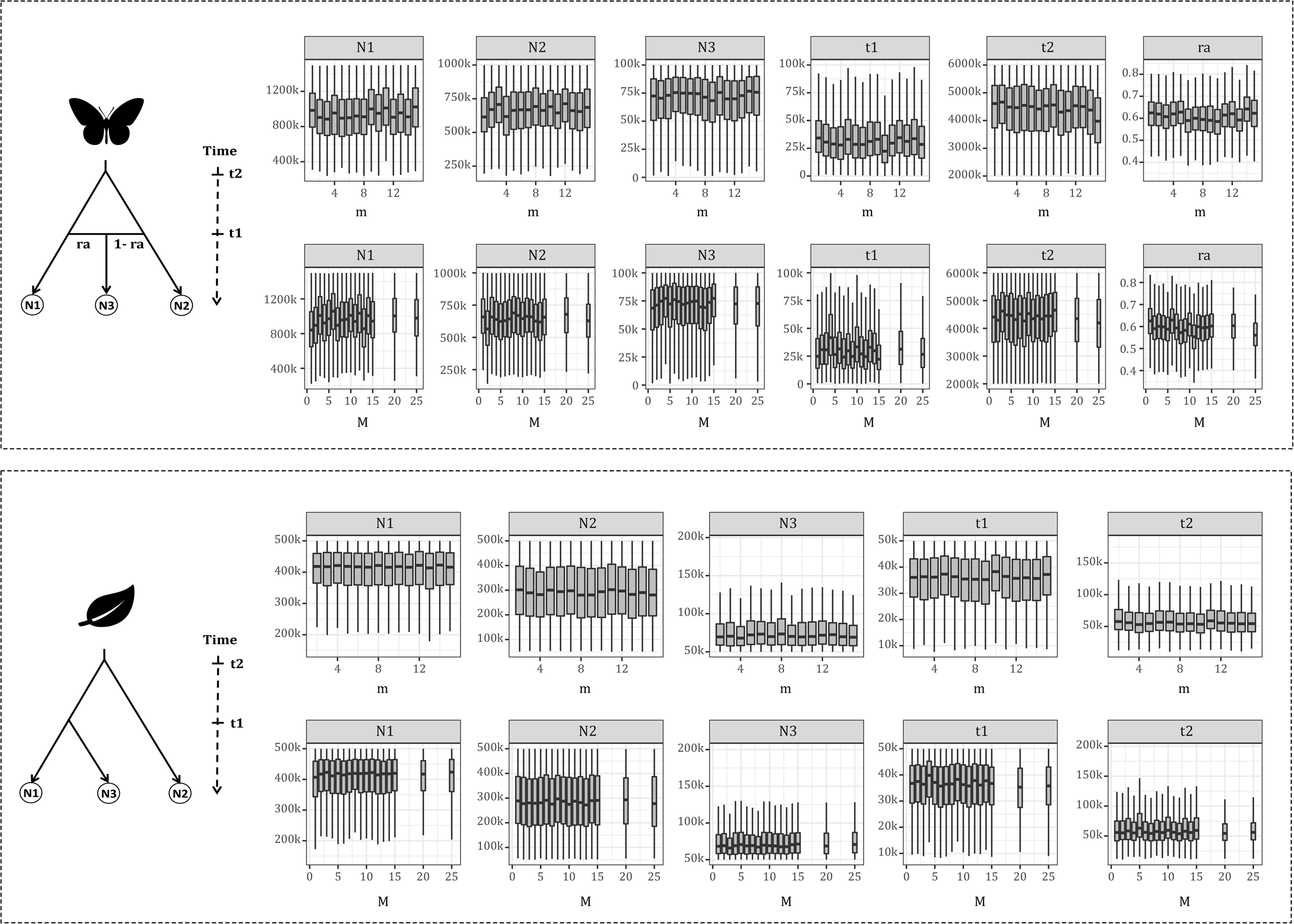
Impact of *ustacks* thresholds **m** and **M** in demographic inferences obtained the ABC procedure. The boxplot summarizes the values of the selected parameters in the 1000 simulations closest to the observed dataset. The parameters include divergence times (t1 and t2) and effective population size (N1, N2, N3) for *Fagus sylvatica* and *Coenonympha* populations, and hybridization contribution (ra) for *Coenonympha*.

## DISCUSSION

Double-digest RAD-sequencing is a widely used technique to investigate population genetics for a wide range of non-model organisms (Peterson et al., 2012; K. R. Andrews et al., 2016). Multiple studies have speculated and tested the impact of different parameter settings on pre- and post-sequencing procedures of RAD-based protocols, especially in loci reconstruction summary statistics, i.e number of loci or SNPs, or heterozygosity (Davey et al., 2013; Gautier et al., 2013; K. Andrews & Luikart, 2014; K. R. Andrews et al., 2014; Puritz et al., 2014; Mastretta-Yanes et al., 2015; Burns et al., 2017; Rochette & Catchen, 2017; Y. Wang et al., 2017; Willis et al., 2017). Nevertheless, only a handful of them linked those parameters to the downstream biological interpretation (Mastretta-Yanes et al., 2015; Rodríguez-Ezpeleta et al., 2016; Shafer et al., 2017; Malinsky et al., 2018). In addition, some pre-sequencing factors, regularly pointed out as sensitive parts of the RADseq protocol (Hohenlohe et al., 2012; Peterson et al., 2012), lack experimental testing on various models that would allow building a solid knowledge of their influence in ddRAD data production. Our objectives here were to test the impact of some wet laboratory and bioinformatic treatment settings on loci recovery as well as on population genetics and demographic inferences.

### Pre-sequencing treatment

Regarding the wet protocol, we focused on two factors for which we did not find any proper evaluation in the literature: the initial DNA quantity and the number of PCR cycles in the last step of library amplification.

The initial amount of DNA required in ddRADseq library preparation is a constraint for small individuals or museum o samples (Blair, Campbell, & Yoder, 2015; Shortt et al., 2017). Here, we found that only a small amount of DNA template (*i.e.* ~50 ng; Fig. 2) was required in our library preparation for the two tested systems, which could open the possibility of using ddRADseq technique with low quantity DNA template from non-invasive sampling in a conservation genomics context. If the initial protocol from Peterson et al. (2012) already suggested to use less than 100 ng of DNA per individual, ddRADseq users commonly process more than 200 ng and even up to 1 μg (Capblancq et al., 2015; Yang et al., 2016; Burns et al., 2017; Sherpa, Rioux, Goindin, et al., 2018). Here we showed that enzyme saturation can already happen at the digestion step with 250 ng. Obviously, the exact amount of DNA that can be digested is directly dependent on the number of enzyme units used and the number of restriction sites. We showed that even a 5-time variation (*i.e.* 250 ng instead of 50 or 150 ng in our study) increased the probability of experiment failure by reducing the loci and the SNP recovery and by inflating the variability between samples (Fig. 2). This points out the need to calibrate finely the amount of DNA and number of enzyme units to avoid a dramatic loss of fragments.

The number of PCR cycles is another part of the experiment that has to be carefully considered (Fig. 2). In their study, Davey et al., 2013 highlighted that PCR cycles introduced GC biases in sequenced RAD libraries. For example, RAD loci with high GC content were sequenced more often compared to RAD loci with low GC content for high numbers of PCR cycles and the opposite was true for for low numbers of PCR cycles (Davey et al., 2013). In this study, our results showed that the number of SNPs and individual heterozygosity is reduced with a low number of PCR cycles, while a high number of PCR cycles increases the number of private alleles. This highlights the trade-off existing between a satisfactory coverage, directly related to the number of PCR cycles, and the limitation of errors occurring during PCR, because these can lead to very weak fragment coverage impeding loci reconstruction (Hohenlohe et al., 2012). Usually the number of PCR cycles is set between 12 and 16 (Peterson et al., 2012; Capblancq et al., 2015; Yang et al., 2016; Burns et al., 2017). Considering the important increase in individual heterozygosity and number of private alleles observed with 25 PCR cycles, our results greatly support this practice. It would be interesting to investigate further if the increase in private alleles and individual heterozygosity could be simply overcome by stringent filtering on allele frequencies aiming at discarding low frequency alleles in the population. Unfortunately, our sampling size did not allow such tests.

Furthermore, we showed that, when properly calibrated, the protocol is greatly reproducible. The triplicates used for 11 individuals of *Fagus sylvatica* showed very close proximities in genetic PCA and most of them showed similar numbers of fragments and coverage (Fig. 3). Such results were consistent with those of Mastretta-Yanes et al. (2015), on which most replicate pairs were clustered together in neighbour-joining dendrograms. Even if there are several steps during ddRADseq laboratory experiment that could lack reproducibility to some extent (*e.g.* digestion/ligation, range size selection, amplification by PCR), our results were robust across replicates. Combined with a low rate of genotyping inconsistency and missing fragments (Fig. 4), our results illustrate that ddRADseq is an accessible method with some key parameters that have to be finely tuned to gain in robustness and reproducibility. Our testing procedure does not claim to cover all parameters that could influence the ddRADseq method but points at key information about lab protocols and gives clues to optimize the technique.

### Post-sequencing treatment

Concerning the dry protocol, an important part of the RAD-based sequencing literature pertains to the bioinformatic treatment of sequences and to loci reconstruction (Mastretta-Yanes et al., 2015; Paris et al., 2017; Rochette & Catchen, 2017; Y. Wang et al., 2017). These studies highlight the impact of clustering thresholds (*e.g.* **M** and **m** parameters of the *STACKS* software procedure) on bioinformatic results and summary statistics. Indeed, these thresholds have been shown to influence the number of recovered loci, coverage, number of identified SNPs and error rates (Mastretta-Yanes et al., 2015; Paris et al., 2017; Rochette & Catchen, 2017; Y. Wang et al., 2017). In agreement with previous works, we found a substantial impact of the minimum coverage (**m**) and clustering (**M**) thresholds on ddRADseq loci recovery and reconstruction during the bioinformatic process. The minimum coverage imposes a minimum number of reads to consider an allele. Many alleles are expected to be lost with a high **m**, while a low **m** can give too much importance to very rare sequences, and thus potentially to sequencing or PCR errors (Catchen et al., 2013). The similarity threshold (**M**) determines the minimum sequence homology to consider that two sequences are variants of the same locus. Choosing a too high **M** can wrongly impede the clustering of different alleles of the same locus, while a too low value could lead to the merging of paralog regions of the genome (Catchen et al., 2013).

We could have expected that such variation in loci reconstruction and SNPs identification would influence, to some extent, the population genetic analyses performed with most ddRADseq datasets. However, our results suggest that the bioinformatic treatment has only a marginal influence on population genetic results. Indeed, no change in genetic differentiation, clustering or demographic inferences were detected neither at the inter-species level for the animal model nor at the intraspecific level for the plant model (Fig 6, 7, S7 and S8).

Moreover, despite a reduced number of individuals, the results of this study are congruent with previous results obtained with a larger sampling for both *Coenonympha* and *Fagus sylvatica* (Capblancq et al., 2015; Capblancq, unpublished data). Such patterns may be explained by the large amount of information generated by the ddRADseq method (10 or 100 of thousands of SNPs). A potential “false” signal due to genotyping inconsistencies at some loci seems negligible compared to the abundance of “true” signal provided by most of the RAD loci. Similar observations have also been made for another species model, *i.e..* Galapagos sea lion (*Zalophus wollebaeki*) in Shafer *et al.* (2017), and similarly, several studies have demonstrated that clustering parameters have no impact on phylogenetic reconstruction (Herrera et al., 2015; Hou et al., 2015; Lee et al., 2018).

Finally, combined with the results from Malinsky et al. (2018) showing that non-random data missingness due to a batch (i.e library) effect had no impact on downstream analyses of the genetic structure. These findings moderate the message from the literature, which commonly presents the bioinformatics treatment as a key parameter of ddRADseq loci reconstruction. If this step indeed has an influence on loci recovery (*i.e* summary statistics), it only has a weak impact on the biological signal resulting from population genetic analyses, at least for the two models tested in this study.

## ACKNOWLEDGEMENTS

The authors belong to the DivAdapt and MALBIO research teams at the Laboratoire d’Écologie Alpine (LECA). We would like to thank Nadir Alvarez and Marianne Gagnon for their useful comments on the manuscript. TCa acknowledges support from the ANR project APPATS (ANR-15-CE02-0004) and all authors acknowledge the LECA for funding support. The LECA is part of the Labex OSUG@2020 (ANR10 LABX56). Most of the computations presented in this paper were performed using the CIMENT infrastructure, which is supported by the Auvergne-Rhône-Alpes region (GRANT CPER0713 CIRA)

## DATA ACCESSIBILITY

All data and scripts will be uploaded on Dryad and publicly accessible after acceptance.

## AUTHOR CONTRIBUTIONS

Thibaut Capblancq, Frédéric Boyer, Tristan Cumer and Charles Pouchon conceived and planned the study. TCa, CP, and Délphine Rioux carried out the lab experiment. TCa and TCu run the majority of the analysis with the help of CP and FB. TCu, CP, FB, Glenn Yannic, Aurélie Bonnin and TCa contributed to the interpretation of the results. TCa and TCu took the lead in writing the manuscript and all authors provided critical feedback and helped shape the analyses and manuscript.

## SUPPLEMENTARY MATERIAL

**Figure S1:** Impact of the bioinformatic threshold **m** (ranging from 1 to 15) on a genetic PCA of *Fagus sylvatica* and *Coenonympha* sp. samples.

**Figure S2:** Impact of the bioinformatic threshold **m** (ranging from 1 to 15) on genetic clustering (sNMF method) of *Fagus sylvatica* and *Coenonympha* sp. samples.

**Figure S3:** Impact of the bioinformatic threshold **m** (ranging from 1 to 15) on Euclidean distances between PCA origin and loci scores in PC1 vs PC2 space, for *Fagus sylvatica* and *Coenonympha* sp. samples.

**Figure S4:** Impact of the bioinformatic threshold **M** (ranging from 1 to 25) on a genetic PCA of *Fagus sylvatica* and *Coenonympha* sp. samples.

**Figure S5:** Impact of the bioinformatic threshold **M** (ranging from 1 to 25) on genetic clustering (sNMF method) of *Fagus sylvatica* and *Coenonympha* sp. samples.

**Figure S6:** Impact of the bioinformatic threshold **M** (ranging from 1 to 25) on Euclidean distances between PCA origin and loci scores in PC1 vs PC2 space, for *Fagus sylvatica* and *Coenonympha* sp. samples.

**Figure S7:** Procrustes superimposition of PCA results for a range of **M** and **m** values and for both the *Coenonympha* and *Fagus sylvatica* models. The two first axes of the PCA were kept to do the Procrustes superimposition among the different **M** and **m** values. The distribution of pairwise correlation coefficients between sets of coordinates resulting from the procruste superimposition are shown for each case.

**Figure S8:** Procrustes superimposition of sNMF results for a range of **M** and **m** r values and for both the *Coenonympha* and *Fagus sylvatica* models. The individual percentages of assignation to the three clusters obtained with sNMF analyses at K = 3 were kept to do the Procrustes superimposition among the different **M** and **m** values. The distribution of pairwise correlation coefficients between sets of assignation scores resulting from the Procrustes superimposition are shown for each case.

**Table S1:** Summary of the samples used in each part of this study.

**Table S2:** Variation of parameters estimation during the ABC procedure for a range of **M** and **m** values and for both the *Coenonympha* and *Fagus sylvatica* models. The minimum and maximum estimation across all **M** or **m** values, and the percentage of variation between them are given for each scenario parameter.

